# Mechanical forces control the valency of the malaria adhesin VAR2CSA by exposing cryptic glycan binding sites

**DOI:** 10.1101/2023.07.31.550984

**Authors:** Rita Roessner, Nicholas Michelarakis, Frauke Gräter, Camilo Aponte-Santamaría

**Affiliations:** Molecular Biomechanics Group, Heidelberg Institute for Theoretical Studies, Heidelberg, Germany; IBMM, University of Montpellier, CNRS, ENSCM, Montpellier, France; Interdisciplinary Center for Scientific Computing, Heidelberg University, Heidelberg, Germany

## Abstract

Plasmodium falciparum (*Pf*) is responsible for the most lethal form of malaria. VAR2CSA is an adhesin protein expressed by this parasite at the membrane of infected erythrocytes for attachment on the placenta, leading to pregnancy-associated malaria. VAR2CSA is a large 355 kDa multidomain protein composed of nine extracellular domains, a transmembrane helix, and an intracellular domain. VAR2CSA binds to Chondroitin Sulphate A (CSA) of the proteoglycan matrix of the placenta. Shear flow, as the one occurring in blood, has been shown to enhance the (VAR2CSA-mediated) adhesion of *Pf* -infected erythrocytes on the CSA-matrix. However, the underlying molecular mechanism governing this enhancement has remained elusive. Here, we address this question by using equilibrium, force-probe, and docking-based molecular dynamics simulations. We subjected the VAR2CSA protein–CSA sugar complex to a force mimicking the elongational tension exerted on this system due to the shear of the flowing blood. We show that upon this force exertion, VAR2CSA undergoes a large opening conformational transition before the CSA sugar chain dissociates from its main binding site. This preferential order of events is caused by the orientation of the molecule during elongation as well as the strong electrostatic attraction of the sugar to the main protein binding site. Upon opening, two additional cryptic CSA binding sites get exposed and a functional dodecameric CSA molecule can be stably accommodated at these force-exposed positions. Thus, our results suggest that mechanical forces, increase the avidity of VAR2CSA, by turning it from a monovalent to a multivalent state. We propose this to be the molecular cause of the observed shear-enhanced adherence. Mechanical control of the valency of VAR2CSA is an intriguing hypothesis that can be tested experimentally and which is of relevance for the understanding of the malaria infection and for the development of anti placental-malaria vaccines targeting VAR2CSA.

## Introduction

*Plasmodium Falciparum* (*Pf*) is responsible for the most virulent and deadliest form of malaria [1, 2]. During the blood stage of the infection, the *Pf* parasite infects, multiplies within, ruptures and reinfects the erythrocytes of the host organism [3]. To achieve this, the parasite substantially modifies the morphology and content of the invaded erythrocytes [4]. In order to avoid the immune response of the host, i.e. the clearance of the infected red blood cells by the spleen, *Pf* sequesters itself in various organs by expressing a multitude of anchor proteins on their surface [5]. This family of proteins, known as Plasmodium falciparum Erythrocyte Membrane Protein 1 (PfEMP1), is encoded by approximately 60 var genes and used to mediate binding on the host’s vascular system [6]. Amongst this adhesin family, VAR2CSA is the sole protein used by the infected erythrocytes for binding to the placental cells through an unusually low-sulphated form of chondroitin sulphate A (CSA) found in the placental intervillous space [7–11] (Figure 1 top).

**Fig 1.**
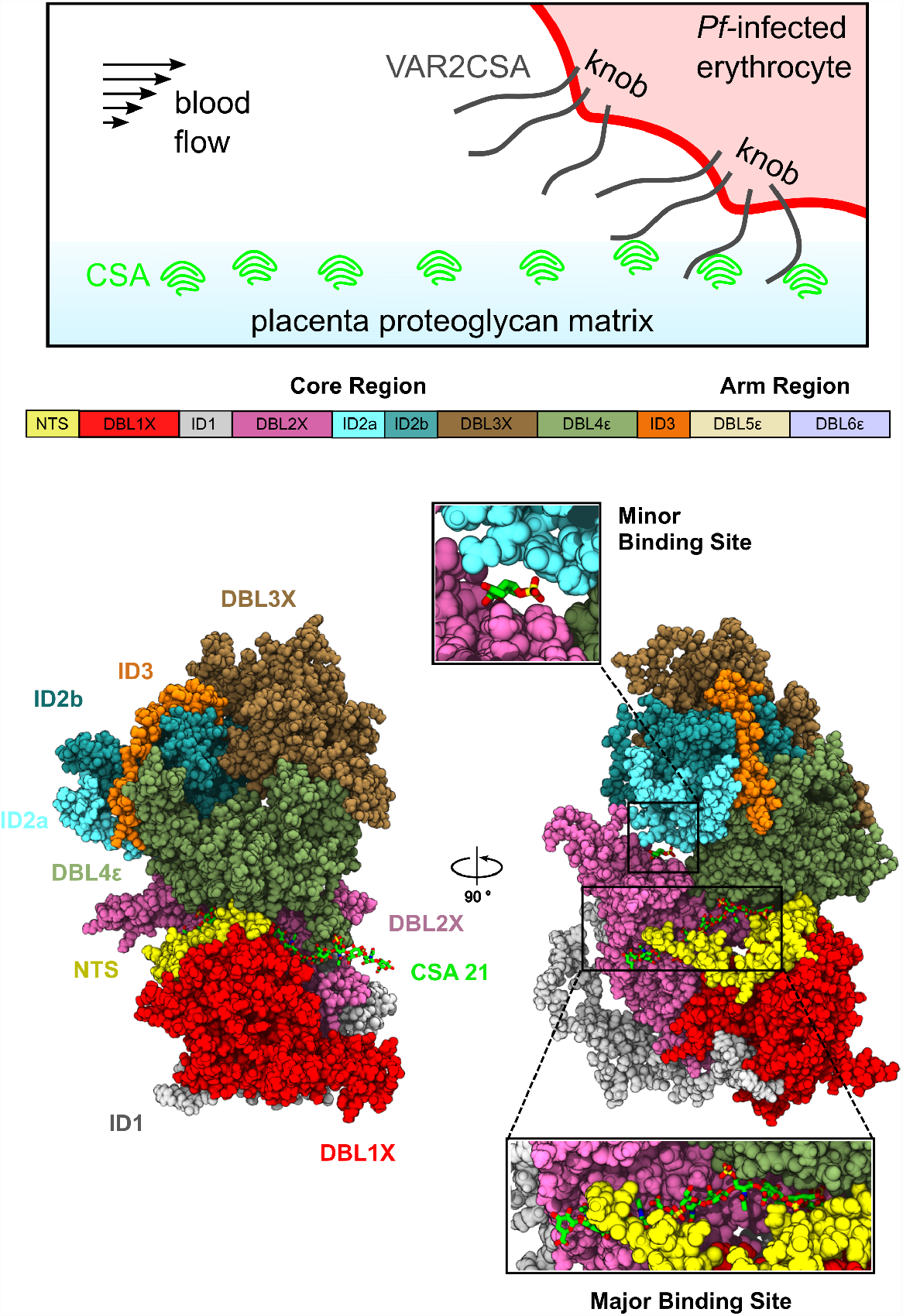
VAR2CSA structure and function. (Top) A schematic depicting the expression of VAR2CSA on a *P. Falciparum* (*Pf*) infected erythrocyte and its attachment to the intervillous CSA, in a blood vessel. (Middle) A schematic of the extracellular core and arm domains of VAR2CSA, colored according to the protein domain. (Bottom) Two views of the VAR2CSA core region (same colours as in the middle panel). The major and minor CSA binding sites are highlighted, with close-up views showing the bound CSA moieties (green sticks).

Pregnancy associated malaria (PAM) affected over eleven million pregnant women in 2020 [2]. Although the infection can be clinically silent, it often leads to maternal anaemia, severely impaired fetal growth, still birth, and spontaneous pregnancy loss. Over 200,000 infant deaths, 10,000 maternal deaths and 900,000 low weight births are attributed to PAM each year alone [7, 12, 13]. Women gain immunity to the disease through successive pregnancies, by acquiring specific antibodies against the VAR2CSA protein [14–16]. Nevertheless, pregnant women are shown to be more susceptible to malaria infection compared to non-pregnant women despite pre-existing immunity [7]. This is attributed to the formation of the placenta which offers a new habitat for *Pf* -infected erythrocytes. Because it is precisely the protein that mediates the adhesion of *Pf* -infected erythrocytes and the CSA proteoglycan matrix of the placenta, VAR2CSA is a promising target for vaccine development against placental malaria [17]. In fact, there are two vaccine candidates at phase I/II clinical trials [18, 19].

VAR2CSA is a multidomain protein composed of extracellular, transmembrane, and intracytoplasmic regions (Figure 1 top). VAR2CSA embeds itself in the membrane of the infected erythrocyte through a single membrane spanning segment. The extracellular region comprises a short N-terminal segment (NTS) and six Duffy-like binding domains (DBL1 - DBL6) which are relatively well conserved amongst parasites [20] (Figure 1, middle). They are connected by four complex, inter-domain regions (ID1 - ID3) which exhibit little homology amongst members of the PfEMP1 family [21, 22] (Figure 1, middle). Due to its large size of approximately 355 kDa, the structural information available until recently was very scarce, relying mostly on the structures of single domains [23–26].

CSA binds with high affinity and specificity to DBL2X and its flanking interdomain regions located at the N-terminal part of VAR2CSA [27, 28]. Structural information of VAR2CSA obtained by low-resolution small angle X-ray scattering (SAXS) revealed the overall shape of the full extracellular region as well as the minimal binding unit [28, 29]. SAXS data from reference [30] revealed that the extracellular region is divided into a compact core and a flexible and potentially-extensible arm (Figure 1, middle). Furthermore, it was suggested that the core region contains two separate CSA binding pores. However, only recently, the cryogenic electron microscopy (cryo-EM) structure of the NF54 VAR2CSA strain was elucidated [31]. The structure contained a CSA dodecamer sugar chain, i.e. the minimal functional binding unit [32, 33], bound to the previously-identified region between the DBL1X, NTS, and DBL2X domains, in the following referred to as the major binding site (Figure 1, bottom). Moreover, this structure displayed additional electron density that was attributed to a single CSA moiety, located in a groove between the DBL2X, ID2a, and DBL4*ϵ* domains, referred to here as the minor or cryptic binding site (Figure 1 bottom). Two apo structures of the VAR2CSA core region (FRC3 strain) [21] displayed high similarity with the holo CSA-bound structure [31]. Furthermore, molecular dynamics (MD) simulations confirmed a similar major CSA-sugar binding site as seen for the NF54 and the FCR3 strain [21]. Taken together, the cryo-EM and simulation data showed that the structural fold of the core region is highly conserved across different VAR2CSA strains and not significantly altered by CSA binding. However, a cryo-EM study of yet another VAR2CSA strain (i.e. 3D7) contradicts these results suggesting large conformational changes upon sugar binding [34].

Cytoadhesion of *Pf* -infected erythrocytes occurs under shear stress caused by the blood flow. Parameters such as the distribution of PfEMP1 on the erythrocyte surface [35] as well as the receptor’s density [36, 37] have been found to be critical for the kinetics of adhesion. For the particular case of VAR2CSA, Rieger and co-workers [36] demonstrated that shear stress enhanced the number of adherent *Pf* -infected erythrocytes expressing this adhesin. This result suggested the possibility of a catch-bond adhesion mechanism for VAR2CSA, in which the protein undergoes a conformational change upon the application of force and thereby increases its adherence. Catch bonds mediate the adhesion of *E. coli* bacteria to mannosylated surfaces, via the force-increased interaction of the fimbrian adhesin protein to its glycan partner [38]. Selectins are another example employing catch bonds during the rolling adhesion of leukocites [39, 40]. Moreover, another PfEMP1 variant has also been suggested to exhibit a catch-bond mode of binding [41]. Catch bonds are contrary to slip bonds in which the binding reduces when they are subjected to force [42].

In this work, we employ equilibrium and force-probe MD simulations to elucidate the, still unknown, molecular mechanism that leads to increased VAR2CSA–CSA adheshion under shear stress. Under the application of elongational force, mimicking the mechanical perturbation VAR2CSA is subjected to by the action of the shear of the flowing blood, we observe a major conformational opening of the VAR2CSA core protein, leading to the exposure of two putative cryptic binding sites prior to dissociation of the tensed CSA chain from the major binding site. We demonstrate that the newly-exposed cryptic binding sites have the ability to stably accommodate a CSA dodecasaccharide. Our simulations explain the preferential opening of the cryptic binding site over sugar dissociation from the major binding site by the orientation of the molecule with respect to the pulling axis and by the strong sugar–protein electrostatic interactions at the major binding site. Overall, our research suggests that rather than increasing the binding affinity of the major CSA binding site, as it would be canonically thought for a catch-bond, force controls the valency of VAR2CSA, switching it from a monovalent to a multivalent state, thereby increasing its adherence to CSA.

## Methods

### VAR2CSA Model Construction and equilibrium MD simulations

The VAR2CSA NF54 strain was considered in this work. A refined and complete model of the core domain (residue IDs 1-1955) was obtained using existing cryo-EM structures of the full VAR2CSA ectodomain (PDB id. 7JGH, 7JGD, and 7JGE [31], and 7B52 [21]), as well as X-ray structures of individual domains (DBL3X: PDB id. 3CML and 3CPZ [24], and 3BQI, 3BQK, and 3BQL [23]) and of the DBL3X-DBL4*ϵ* complex (PDB id. 4P1T [26]). The structures were combined into a modelling protocol with Modeller 9.18 [43] to complete missing residues and loops that were absent in the full ectodomain structures. The modelled segments comprising residues 407–485 and 529-556 were rotated manually after the modelling procedure to prevent their wrapping around the protein. The CSA dodecamer found in the major CSA binding site of the NF54 structure was extended to 21 sub-units using PyMOL. Subsequently, the system was energy minimized with the steepest descent algorithm applying harmonic restraints on the atoms which were observed in the NF54 structure (harmonic elastic constant was 5000 kJ/mol/nm^2^ and reference positions for the harmonic potentials were taken from the NF54 [31] structure). The resulting model displayed a root mean square deviation (RMSD) of 0.23 Å from NF54 structure (PDB id. 7JGH [31]) and of 1.43 Å from the FCR3 strain structure (PDB id. 7B52 [21]). The final model was then equilibrated through the use of equilibrium MD simulations, following a protocol of progressive restraint relaxation as explained as follows.

After energy minimization, the conformation of the loops (residues 407–485 and 529–556) was refined with Langevin stochastic dynamics in vacuum for 100 ns. The rest of the system was kept frozen at a temperature of 500 K with a friction time constant of 0.002 ps. The protein was subsequently solvated with water and 150 mM NaCl in a dodecahedron box with an excess of chloride ions to neutralize the net charge of the protein. The resulting system consisted of approximately 0.7 M atoms. The solvated system was energy minimized and simulated under NVT-ensemble conditions for 500 ps at a temperature of 310 K. All heavy atoms were position-restrained with a force constant of 1000 kJ/mol/nm^2^. In a second stage, the system was equilibrated in an NPT ensemble, under the same conditions, for 1 ns. This step was repeated with all non-modelled atoms being restrained with a force constant of 1000 kJ/mol/nm^2^. The system was further equilibrated in an NPT ensemble for 1 ns with position restraints on the backbone atoms. The force constant in this instant was 100 kJ/mol/nm^2^. Finally, the system was simulated under the same conditions for 200 ns, without any backbone restraints. Ten replicas were simulated for a total of 2 *μ*s of cumulative simulation time.

### Force Probe MD Simulations

Two frames of each replica from the equilibrium simulations described above were extracted and used as starting points for 20 pulling simulations (one frame was taken at the end of the simulation and a second frame 30 ns before). Each conformation was subsequently re-solvated in a cubic box with water and ions (as the simulations above) resulting in a system of approximately 1 M atoms. The solvated systems were independently thermalized via a 500 ps NVT simulation, maintaining protein backbone atoms position-restrained with a force constant of 100 kJ/mol/nm^2^. The force application points were selected to mimic the elongational tension experienced by the core domain of VAR2CSA under blood flow, namely based on the C-terminal anchoring of VAR2CSA in the erythrocyte membrane and the 1’-O-glycosidic linkage of gluceronic acid to the placental proteoglycans [44]. Accordingly, the first residue of the bound CSA molecule (numbering according to Figure 2A) and the C-terminal methionine residue of the core protein were subjected to harmonic spring potentials with a force constant of 1000 kJ/mol/nm^2^ moving with a constant velocity of 0.1 m/s in opposite directions. Each replica was simulated for 100 ns for a cumulative force-probe simulation time of 2 *μ*s.

**Fig 2.**
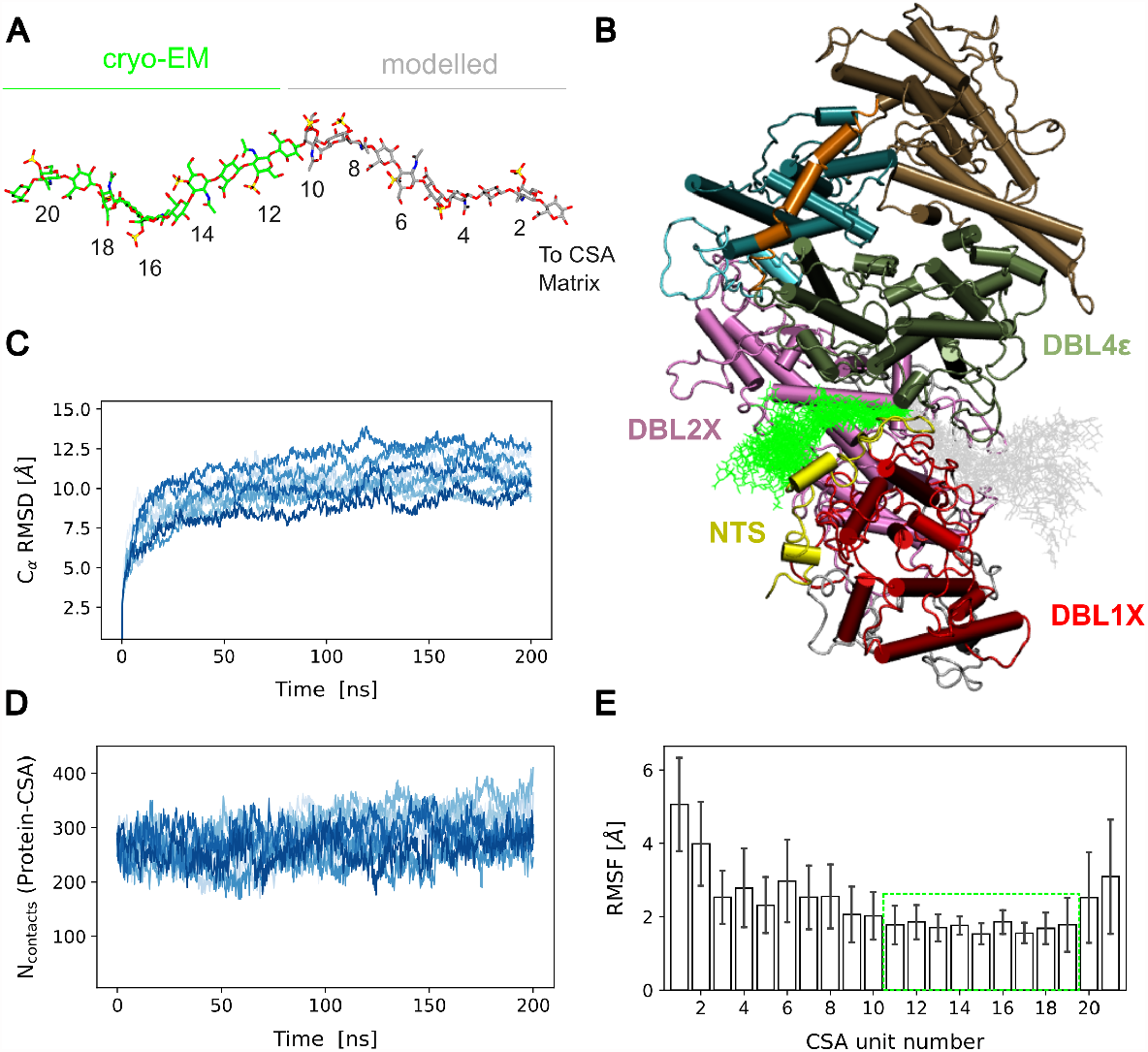
MD Equilibration of VAR2CSA with a CSA 21mer sugar chain bound to it. (A) The CSA molecule is presented in stick representation (green: cryo-EM-resolved part, PDB id. 7JGH [31]; grey: modelled part). The numbering of the monosaccharides increases with respect to the terminus that points to the CSA matrix. (B) Example of the conformations sampled by the CSA 21-mer in one of the 200 ns MD simulations when the molecule was bound to the major binding site of the core region of VAR2CSA. The sugar is shown as sticks (colored as in A) and the protein is displayed in cartoon representation (colored according to Figure 1). (C) C*α* root mean square deviation (RMSD) from the initial conformation of the VAR2CSA core for n=10 independent replicas (differently colored lines). (D) Time traces of the number of contacts between VAR2CSA and the CSA chain bound to the major binding site (n=10 independent replicas). (E) Root mean square fluctuation (RMSF) of each CSA monosacharide calculated over the last 100 ns of the n=10 simulation repeats (average ± standard deviation, n=10). Green Box highlights the part of the sugar resolved by cryo-EM, which exhibited a relatively low conformational flexibility.

### CSA dodecamer Docking at cryptic binding sites and MD relaxation

To understand whether force-exposed cryptic binding sites have the ability to accommodate a functional CSA molecule, we docked a CSA dodecamer to the open VAR2CSA molecule using the HADDOCK webserver [45]. We selected this size of saccharide since a dodecamer is the minimum length needed for CSA binding in the major CSA binding site [32, 33]. The dodecamer was treated as “ligand” while the VAR2CSA together with the 21mer sugar chain bound to the major binding site were considered as the “receptor” during the docking calculation. The structure of the dodecamer (ligand) was taken from the cryo-EM structure (PDB id. 7JGH [31]). For the protein with the bound sugar 21mer (receptor), we selected an open conformation with the N-terminal and C-terminal subregions separated by a distance of 9 nm (see Figure 3D) from each replica of the pulling simulations, resulting in 20 initial input receptor configurations. The dodecamer was docked at two different locations of the protein: one corresponding to the minor binding site, which locates at the interface between the DBL2X and the ID2a domain, and the other corresponding to the DBL4*ϵ* domain portion of the major binding site that got exposed upon application of force. Accordingly, based on the cryo-EM holo structure of VAR2CSA (PDB id. 7JGH [31]), the active residues directly involved in the interaction were assumed to be T910, Y911, T912, T913, H949, K952, D968, and K970 for the minor binding site and H1782, I1783, G1784, I1785, I1878, M1879, E1880, K1889, R1890, N1896, N1898, and Y1899 for the DBL4*ϵ* site. In addition, the first five sugars of the dodecamer were treated as the active region involved in the interaction. No passive residues were considered in the docking calculation. The number of structures for rigid body docking was set to 10000 while the number of structures for both semi-flexible refinement and explicit solvent refinement was 400. Default values were considered for the all the other docking parameters. HADDOCK clustered the docking conformations into into eight different groups. A pose representative of each cluster was selected. Each docked conformation was subsequently solvated and equilibrated in the same manner as for the force-probe simulations. The stability of the docked dodecamer was monitored in 100 ns simulations. To avoid re-closing of the VAR2CSA sub-regions, we applied a weak flat-bottom potential to the first residue of the CSA molecule and the C-terminal residue of the core protein, which was not zero when their distance got below a distance of 13 nm. The “flat-bottom-high” pulling coordinate option in GROMACS was used for this purpose. The elastic constant of this potential was 1000 kJ/mol/nm^2^. 100 ns times 8 poses times equals 800 ns of cumulative simulation time per binding site.

**Fig 3.**
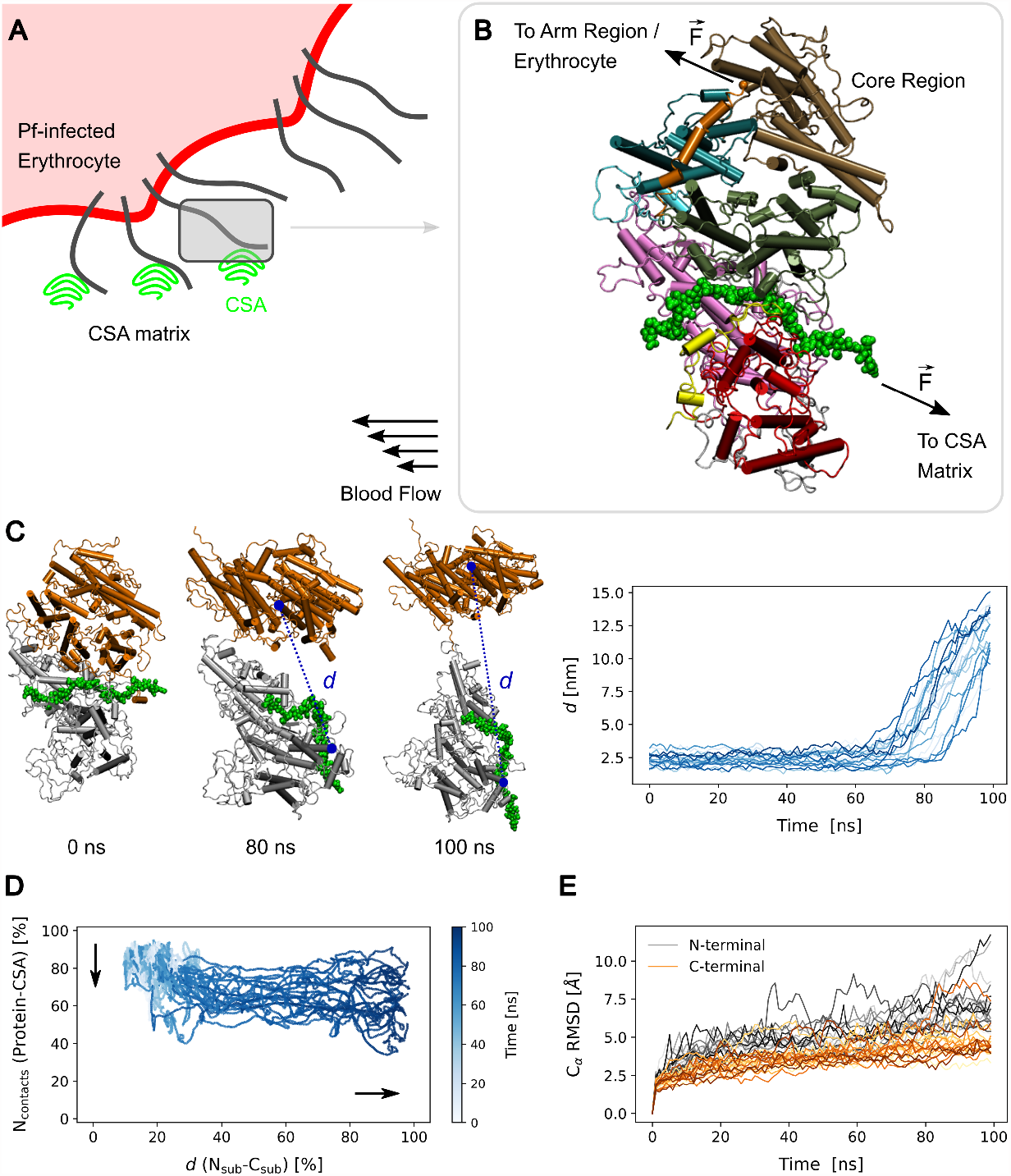
Force-induced opening of the VAR2CSA core region. (A) A schematic depicting the adhesion of a *Pf* -infected erytrhocyte (red) to CSA molecules of the intervillous matrix (green), mediated by VAR2CSA (black lines). The shear of flowing blood drags the erythrocyte. In consequence, elongational force acts on VAR2CSA. (B) To mimic this mechanical stress, forces 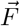 were exerted on the VAR2CSA core–CSA complex during pulling simulations (VAR2CSA core: cartoon, colored according to Figure 1, CSA: green spheres). Force was applied to the first monomer of the CSA chain (transmitting the tension from the anchor point at the CSA matrix) and at the C-terminal methionine residue of the core region (linked to the infected erythrocyte by the VAR2CSA arm region). (C) Opening of the VAR2CSA core region was monitored during the course of the pulling simulations by measuring the separation distance *d* (dashed blue line) between the residues 120 and 1663. Left: The N-terminal sub-region (domains NTS, DBL1X, ID1 and DBL2X, i.e. residues 1–964) is colored in gray while the C-terminal one (ID2a, ID2b, DBL3X, DBL4*ϵ*, and ID3, i.e. 965–1954) is shown in orange. The CSA moiety (green spheres) is shown to highlight the area of the major binding site and its movement during the simulation. Right: *d* time traces for all 20 independent simulation replicas. See also movie S1. (D) The number of VAR2CSA–CSA contacts (y-axis) is presented as a function of the distance between the N-terminal- and C-terminal sub-region (x-axis) and as a function of time (color) for the 20 simulation replicas. The earlier inversely relates to the dissociation of the CSA 21mer from the major binding site (vertical arrow) whereas the latter corresponds to opening of the interface between subregions (horizontal arrow). (E) C*α* root mean square deviation (RMSD) of the N-terminal-(grey) and C-terminal sub-region (orange) as a function of time.

### Simulation algorithms and parameters

Through all the simulations described in this work the temperature was maintained at 310K through the use of the Nose-Hoover thermostat [46, 47], coupling the protein and the rest of the system separately to the thermostat, and using a coupling time constant of 1 ps. The pressure was kept constant at 1 atm by means of the Parrinello-Rahman barostat [48, 49] (coupling time constant of 5 ps). The GROMACS 2019.4 [50] molecular dynamics package was used for the simulations with an integration time step of 2 fs. The CHARMM36m forcefield [51] was used for the protein, the CHARMM-modified TIP3 model [52] for the water molecules, and CHARMM parameters for the CSA sugar and the NaCl ions. The CHARMM-GUI [53–55] web-server was used for the generation of the sugar forcefield parameters and the GROMACS gmx tools for the remaining force-field terms. Electrostatic interactions were calculated by using the Particle Mesh Ewald method [56, 57] in the direct space within a cut-off distance of 1.2 nm and in the reciprocal space beyond that cut-off. Electrostatics were treated with reaction-field in the in-vacuum loop relaxation step with a dielectric constant of 1. Short-range interactions were modelled through a Lennard Jones potential within a cutoff distance of 1.2 nm. Neighbor atoms were considered through the Verlet buffer scheme [58]. The Lennard Jones force was shifted between 1 nm and 1.2 nm. Bonds involving hydrogen atoms were constrained by using the LINCS algorithm [59] and both bonds and angles of water molecules were constrained via SETTLE [60].

### Simulation analysis

The simulation analysis was performed with GROMACS tools [50] and MDAnalysis 2.3 [61, 62]. Root-mean-square deviations (RMSD) of atomic distances and root-mean-square fluctuations (RMSF) of atomic positions were determined with the MDAnalysis.analysis.rms module [63, 64].

#### Contact Analysis

Contact analysis was performed using the GROMACS tool *gmx mindist*. To evaluate the stability of the VAR2CSA/CSA complex in the equilibrium MD simulations, we computed the number of contacts between the protein and CSA over the course of the trajectories (Figure 2D). We monitored the opening of the VAR2CSA core in the force probe simulations by calculating the number of VAR2CSA—CSA contacts as a function of the distance between the N-terminal- and C-terminal sub-region and as a function of time (Figure 3D). We also computed the number of contacts between the CSA dodecamer docked to the DBL4*ϵ*- and DBL2X cryptic binding site to evaluate its interaction with the VAR2CSA/CSA complex (Figure 5C).

**Fig 4.**
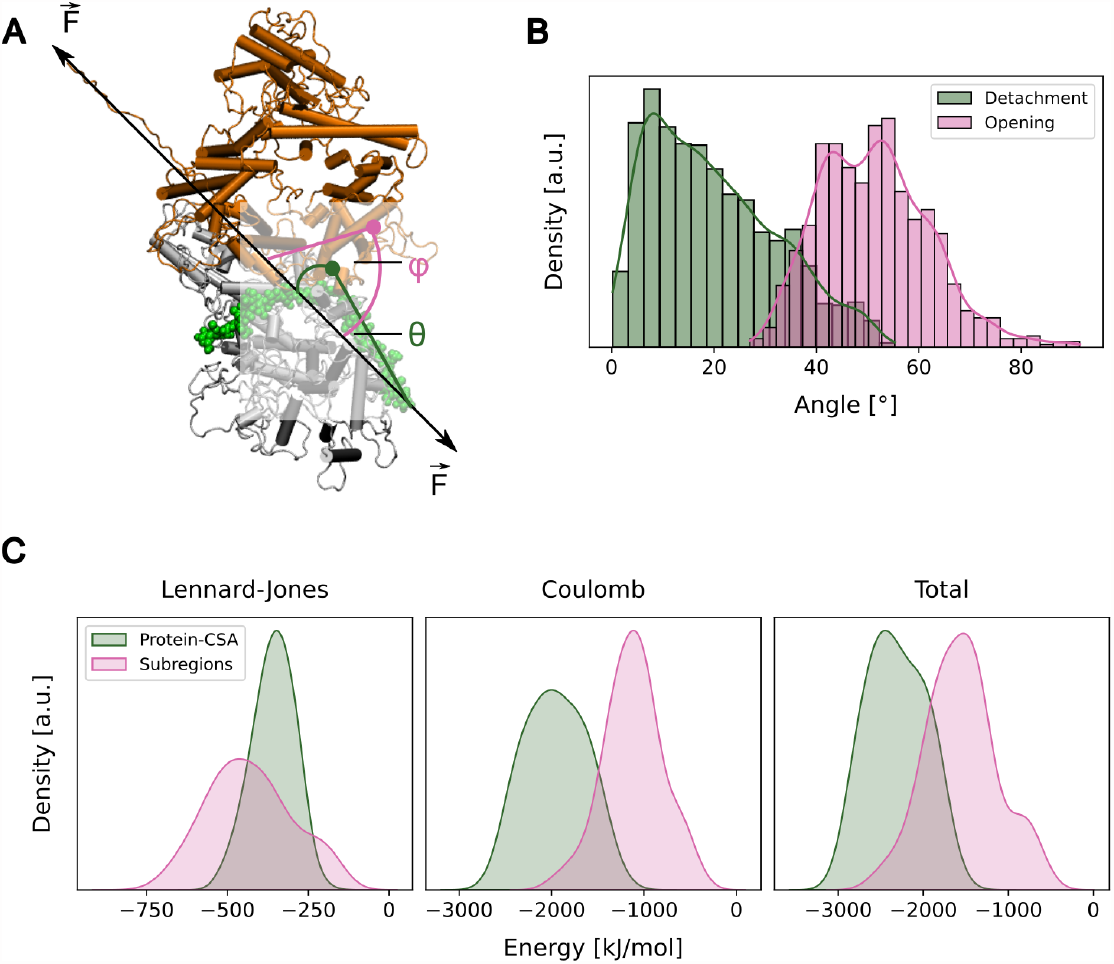
Pulling orientation of VAR2CSA and protein–sugar electrostatic interactions can explain the opening of the tethered VAR2CSA core region. (A) Cartoon showing the two angles, *θ* and *ϕ*, used for the exploration of the CSA detachment and opening mechanisms, respectively. *θ* is the angle formed by the CSA chain bound at the major binding site with respect to the pulling axis (green). *ϕ* is the angle along the sub-region interface with respect to the pulling axis (pink) (B) Histogram showing the distribution of *θ* (green) and *ϕ* (pink) angles calculated over the 20 force-probe simulation repeats, until the sub-regions were entirely separated. (C) The potential energy between the sub-regions that opened under force (sub-region–sub-region: pink) and between the CSA chain and the major binding site (protein–CSA: green) were computed from the equilibrium MD simulations. Normalized distributions of the short-range Lennard-Jones (left) and Coulomb (middle) contributions, as well as the distribution of the sum of them (right), are presented.

**Fig 5.**
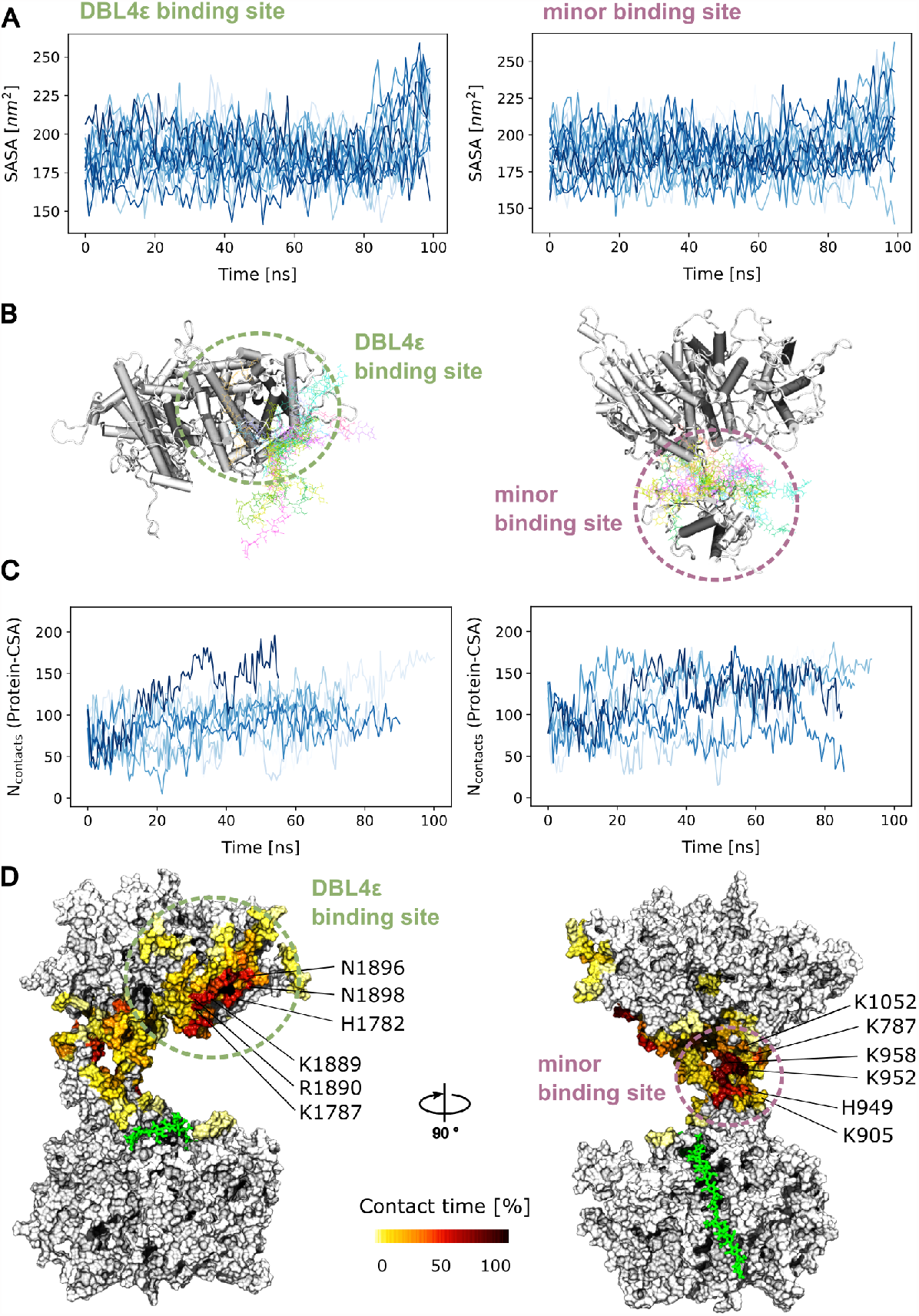
Force exposes two cryptic secondary CSA sugar binding sites. (A) Time-traces of the sugar accessible surface area of the DBL4*ϵ* portion of the major binding site and of the minor binding site, located at the interface between the DBL2X, ID2a, and DBL4*ϵ* domains, extracted from the pulling simulations (n=20 for each case). Examples of the conformations adopted by a CSA sugar dodecamer when it was docked and relaxed (by MD simulations) at these two sites (left: DBL4*ϵ*- and right: minor binding site) (sugar shown as colored sticks and protein in cartoon). For clarity, only the relevant portion of the core protein was shown. (C) Time-traces of the number of contacts established by the docked dodecamer and the protein during the MD relaxation simulations (n=10; left: DBL4*ϵ*- and right: minor binding site). (D) Contact time between protein residues at each identified binding site and the docked dodecamer were recovered from the MD relaxation simulations. The contact time is color-coded on the surface of the protein, relative to the total number time of the simulations (n=8 amounting for 800 ns of cumulative simulation time). Positively-charged residues displaying a large contact time are indicated. The sugar 21mer that remained bound to the DBL2X part of the major binding site is shown as green sticks.

#### Opening of VAR2CSA core

The opening of the VAR2CSA core was additionally monitored over the course of the force probe simulations by measuring the distance between the residues 120 and 1663 of the N-terminal- and C-terminal sub-region, respectively (Figure 3C).

#### Orientation analysis

To shed light on the mechanism of CSA detachment and VAR2CSA opening, we evaluated the orientation CSA bound to the major binding site as well as the orientation of the sub-region interface with respect to the pulling axis over the course of the force-probe MD simulations (Figure 4B). The vector between the points of force application on the VAR2CSA and the pulled CSA molecule defined the pulling direction. A suitable third point, consisting of CSA-residue 11 and of protein residue F1744, defined the orientation angles *ϕ* (CSA detachment) and *θ* (VAR2CSA opening), respectively (Figure 4A).

#### Potential energy of VAR2CSA–CSA and subregion interaction

The short-range Lennard-Jones- and Coloumb contributions of the interaction potential between VAR2CSA and CSA as well as between the N-terminal- and C-terminal sub-region were computed using the GROMACS *gmx mdrun* utility (Figure 4C).

#### Solvent accessible surface area (SASA) at cryptic binding sites

We computed the SASA of the DBL4*ϵ* binding site (residues N1782, K1787, K1889, R1890, N1896, N1898) and the minor binding site (residues K797, K905, H949, K952, K958, K1052) using the GROMACS *gmx sasa* over the course of the force probe simulation to monitor their force-induced exposure (Figure 5A). To take into account the size of the sugar chain, a solvent probe radius of 2.0 nm was used for this calculation.

The molecular visualization package VMD [65] was used to render the snapshots of the system as described above.

## Results

### A refined model of the VAR2CSA core region in complex with a CSA dodecamer

We first generated a complete model of the VAR2CSA NF54 strain core region (ranging from the NTS to the ID3 domain) in complex with a CSA 21mer bound to the major binding site (Figure 1). The model is based (mainly) on the recently determined cryo-EM structures of this strain [31] and the VAR2CSA FRC3 strain [21]. We considered the CSA dodecamer that was observed in the cryo-EM structure and prolonged it by 10 monomers, to be able later to properly examine the force-response of the protein–sugar complex (Figure 2A). We then relaxed the model by conducting n=10 independent equilibrium MD simulations of 200 ns each, for a cumulative simulation time of 2 μs (Figure 2B). Over the course of the simulations, the system stabilized within 150 ns at conformations that deviated by no more than 15 Å (carbon-alpha root mean square deviation, RMSD) from the initial conformation (Figure 2B,C). We find these are reasonable RMSDs given the large size of the protein and the abundance of flexible loops. Importantly, the number of contacts between the CSA sugar and the VAR2CSA protein fluctuated around a constant value, indicating that the sugar stayed stably-bound to the protein over the course of the simulations (Figure 2D). Each monosaccharide displayed structural fluctuations of less than 5 Å(Figure 2E), which are similar to the fluctuations observed in a previous study of the FCR3 strain [21]. As expected, the part of the sugar that was observed in the cryo-EM structure displayed slightly lower structural fluctuations than the modelled part. Consequently, the MD structural relaxation generated a refined ensemble of conformations that allowed us to investigate the mechanical response of the VAR2CSA-CSA complex.

### Force opens the core region of tethered VAR2CSA

In order to understand the force-response of the VAR2CSA-CSA complex, we performed force-probe MD simulations, applying force in a manner that mimics the elongational tension the protein is subjected to, when it adheres a *Pf* -infected erythrocyte to the CSA proteoglycan matrix under the action of a shear flow (Figure 3A). Under these conditions, VAR2CSA is expected to be pulled away from the tethering CSA sugar chain by the sheared *Pf* -infected erythrocyte. We considered the core region of the protein which contains the main CSA binding sites (Figures 1 and 3B). The “arm” region, consisting of domains DBL5*ϵ* and DBL6*ϵ*, is not in direct contact with the CSA sugars [21, 30, 31] (Figure 3B). Accordingly, we applied a harmonic force on the C-terminal methionine residue of the core region (residue id. 1953), assuming that force is transmitted there by the arm region. An opposing harmonic force was applied to the first monosaccharide of the bound CSA 21mer, i.e. the point that tethers VAR2CSA to CSA (Figure 3B). The two pulled points were separated at a constant speed of 0.2 m/s. We extracted 20 conformations from the equilibrium MD simulations of the VAR2CSA core–CSA complex and used them as starting configurations for 20 independent force-probe MD simulations. Upon application of force, we observed a substantial opening of the core region: the N-terminal sub-region composed of the domains NTS, DBL1X, ID1 and DBL2X (shown in gray) shifted away from the C-terminal sub-region constituted by ID2a, ID2b, DBL3X, DBL4*ϵ*, and ID3 (depicted in orange) (Figure 3C). Another possibility could have been that force induced the detachment of the sugar from the major binding site before the core region opened. This possibility was ruled out after plotting the the number of contacts between the protein and the sugar versus the distance between regions (Figure 3D). As the simulation time progressed, we observed that the system followed preferential paths in which opening always occurred first (see distance between sub-regions decrease to a larger extent than protein–sugar contacts in figure 3D). Note that the fold of these two sub-regions was largely preserved under force application, apart from a residual unfolding of the loose C-terminus (Figure 3E). This is mainly attributed to the large number of disulfide bonds that helped VAR2CSA maintaining its structural integrity. Thus, elongational tension opens the core region of tethered VAR2CSA and separates it into two almost structurally-intact sub-regions.

### Orientation of VAR2CSA and sugar–protein electrostatic interactions explain VAR2CSA opening preference over sugar dissociation

We next shed light on the mechanism behind the preferential opening of the core domain prior to detachment of the sugar from the major binding site. Because the pulling direction influences the force response of biomolecules [66], we first checked the effect of the orientation of VAR2CSA with respect to the pulling axis. We calculated the angles formed by the tethered sugar and the opened interface with respect to the pulling axis, denoted as *θ* and *ϕ*, respectively (Figure 4A). We observe that the CSA sugar inclined with respect to the pulling axes forming a *θ* angle of 20 ± 12 ^*o*^ (average ± standard deviation) (Figure 4B). The interface between the N- and the C-terminal sub-regions, in contrast, adopted a more perpendicular orientation with respect to the pulling axis (*ϕ* = 51 ±11 ^*o*^, average± standard deviation, Figure 4B). These numbers suggest that the orientation of the molecule along the line of tension causes the interface of the cryptic binding site to open in “zipping” mode whereas the detachment of the CSA molecule from the major binding site follows a mechanically more resistant “shearing” motion.

In addition to the orientation, we checked whether the interaction strength between the interfaces subjected to force had an influence on the opening behavior. To achieve this, we extracted the potential interaction energy between the CSA chain and the major binding site from the equilibrium MD simulations and between the two sub-regions (Figure 4C). Note that the potential energies only allow an approximate relative analysis on the energetics. Free energies would be required for a full quantitative picture. The difference in contact area resulted in a broader and overall stronger short-range Lennard-Jones contribution to the interaction between the sub-regions compared to the CSA-VAR2CSA interaction (Figure 4C, left). The strong electrostatic interaction of the (negatively-charged) CSA sugar with the (positively-charged) major binding site was notorious in the Coulomb contribution compared to the sub-region interface (Figure 4C, middle). Altogether, the potential interaction energy between the CSA sugar and the major binding site was stronger than that between subregions, mainly due to the electrostatic contributions (Figure 4C, right). Thus, the sugar–protein electrostatic interactions also contribute to the opening of the interface while the CSA chain remains attached to the major binding site.

### Force causes the exposure of two cryptic CSA binding sites

We finally investigated the functional consequences of the opening of the VAR2CSA core due to the application of force. Remarkably, the opening of the core region detached the DBL4*ϵ* from the DBL2X domain, thereby increasing the level of exposure of the DBL4*ϵ* residues that were engaged in direct interactions with the sugar chain at the major binding site (Figure 5A, left). Note that despite of the loss of such interactions, during the force-induced opening process, the CSA 21mer remained bound to the major binding site, more specifically to the DBL2X part of it (Figure 3D). In addition, the minor binding site at the groove of the DBL2X, ID2a, and DBL4*ϵ* domains increased its accessibility upon opening (Figure 5A, right). The exposure in this case was less notorious than for the DBL4*ϵ* site, because this minor binding site was already partly exposed before the application of force.

A CSA chain could potentially be accommodated at these two newly-exposed sites. We tested this hypothesis by docking a CSA dodecamer, the minimal functional binding unit [32, 33], separately in either of such positions. We allowed the docked structures to relax in simulations of several tens of ns (from 55 to 100, ns) while keeping the protein in the open state (see methods). During this process, the CSA molecule remained stably bound at either of these two sites (Figure 5B). A quantitative indication of this is the stable fluctuations around a constant value in the number of CSA sugar–protein contacts, i.e. no loss of such contacts over time (Figure 5C). Due to its electrostatic nature, these contacts occurred often between the negatively-charged moeities of the CSA chain and positively-charged residues of of the protein (Figure 5D). This result suggests that, apart from the major binding site, VAR2CSA features two other sites where a CSA dodecamers can stably bind. These two sites are cryptic but get exposed upon opening of the core region of the VAR2CSA, due to the application of an external elongational force.

## Discussion

In this work, we shed light on the molecular mechanism governing the increased VAR2CSA–CSA interaction under shear stress using equilibrium and force probe MD simulations.

Our data demonstrate that upon force exertion, the core region of VAR2CSA undergoes a major conformational rearrangement by opening up two structurally-intact sub-regions (Figure 3). The CSA molecule already bound at the major binding site operates as an anchor point, allowing for the opening of VAR2CSA to occur. This causes the subsequent exposure of two cryptic binding sites, which have the ability to accommodate further CSA molecules, prior the dissociation of CSA from the major binding site (Figure 5). This is a remarkable result, as it proves that force does not cause neither the immediate dissociation of CSA from VAR2CSA nor its unfolding, but instead it prepares the adhesin to bind further CSA chains (or another parts of the same chain) at two other locations. Consequently, our data support a mechanism in which application of mechanical forces on tethered VAR2CSA increase the valency of this protein to CSA, and with it its avidity, to potentially reinforce its attachment to the proteoglycan matrix.

The recent low-[30] and high-resolution [21, 31, 34] structural information of VAR2CSA clarified the exact location of the minimal functional CSA binding unit around the DBL2X domain [27, 28] (i.e. the major binding site), but also suggested the possibility of a second CSA binding location, indicated by a single ASG monosaccharide observed in the holo cryo-EM structure of VAR2CSA [31](Figure 1). One of the cryptic binding site exposed by force in our simulations precisely coincides with this proposed additional (minor) binding site (Figure 5). The other proposed binding site is precisely the region of the DBL4*ϵ* domain constitutive of the major binding site. When detached from the DBL2X domain, upon opening, this region had the ability to stably accommodate a dodecameric CSA chain by itself, while the protein was held by another sugar at the major binding site. Taken together, this reveals a key role of these two regions, namely the activation of additional CSA binding sites. Force, accordingly, controls the level of exposure of these sites.

Because of its dimensions (∼17 nm in the compact state), VAR2CSA is suggested not to be directly influenced by the hydrodynamic shear of the flowing blood. Instead, VAR2CSA is assumed to experience an elongational tension when it is tethered to a CSA chain, at one side, while it is pulled by the infected erythrocyte by the action of the shear flow, at another side. Consequently, we applied a force along VAR2CSA that mimics such tension (Figures 3A,B). This resembles the behavior of the blood clotting protein von Willebrand factor, which also becomes mechano-activated by the influence of an in-homogeneous tension distribution along the molecule generated by the shear of flowing blood [67, 68]. Although our choice is a reasonable assumption, in the future, additional simulations at a lower level of resolution but explicitly imposing a shear flow [69] would be required to establish the connection between elongational tension acting on VAR2CSA and the shear flow acting on the CSA-VAR2CSA-erythrocyte system.

We also investigated the mechanism leading to the conformational opening of VAR2CSA prior to the sugar dissociation from the major binding site (Figure 4). A strong influence of the pulling direction on the force response of biomolecules has been previously observed [66, 70]. A similar effect applies here for VAR2CSA. The angles of the CSA–major binding site interface and the angle of the subregion–subregion interface with respect to the force vector suggest that they open through a “shearing” and “zipping” motion, respectively (Figures 4A,B). In competition, shearing of the sugar from the major binding site would be less likely to occur than the unzipping of the subregion interface. In addition, the interaction energy between the CSA chain and the major binding site was found to be stronger than that between the subregions, mainly due to the strong electrostatic attraction between the negatively-charged CSA moieties and the positively charged residues of the binding site (Figure 4C). Accordingly, the VAR2CSA–sugar interactions are highly force resistant due to, first, their orientation parallel to the force axis, which leads to a shearing mode for dissociation as opposed to the unzipping of the VAR2CSA core-subregions, and, second, the strong charge complementarity between the sugar and the protein major binding site. This dictates the preference for the opening and subsequent activation of VAR2CSA over its dissociation and with it that of an infected erythrocyte from the CSA sugar.

The observation of shear-enhanced adhesion for VAR2CSA [36] suggested that this adhesin employs a catch-bond mechanism, i.e. its adherence increases upon the application of force [42]. Such behavior is utilized by bacteria to attach to their glycan substrate [38] or by rolling leukocytes [39, 40]. It has also been proposed for another PfEMP1 variant [41]. However, locally, force did not seem to engage the CSA chain in a stronger interaction with the major binding site, as indicated by a loss of sugar–protein contacts upon force application (Figure 3D). Accordingly, we propose the major binding site of VAR2CSA does not strictly follow a catch-bond behavior. Instead, force exposed two other sugar binding sites, which were cryptic under equilibrium conditions and where a CSA glycan could be stably maintained (Figures 5). Thus, VAR2CSA switches from a monovalent to a multivalent state by the action of force, thereby amplifying its avidity for CSA binding. Multivalency is a general mechanism to enhance ligand-receptor interactions [71] and a central feature for protein-glycan recognition [72]. We propose that VAR2CSA follows this principle to enhance its adhesion to the placental CSA matrix, but here controlled by the elongational tension acting on the protein as a result of the shear of flowing blood.

Our data emphasize the importance of conducting shear-binding assays [36], and not only static ones, to study VAR2CSA adherence. More specifically, our computational prediction of force control of the valency of VAR2CSA could be tested in such type of experiments. For instance, the opening of the core region could be artificially prevented, by introducing cross-linking cysteine residues at the N- and C-terminal subregions, and with this the shear-enhanced response is expected to be abrogated.

## Conclusions

Based on equilibrium and force-probe MD simulations, we propose a mechanism for the interaction of the *Plasmodium falciparum* malaria adhesin VAR2CSA with the placental CSA matrix. In this mechanism, mechanical force modulates the valency of tethered VAR2CSA by opening of the core in two structurally-folded regions. Opening causes the exposure of two cryptic CSA binding sites. This enables VAR2CSA to stably bind a second CSA dodecamer in addition to the one tethered to the major binding site. We attribute the preferential opening of the protein (and consequent exposure of cryptic binding sites), prior to the dissociation of CSA from the major binding site, to the orientation VAR2CSA adopts under tension and to the strong sugar–protein electrostatic interactions occurring at the major binding site. Rather than a catch-bond influencing the affinity of the major binding site, control of the valency of VAR2CSA by force is a possible explanation of the observed shear-enhanced adherence of *Pf* -infected erythrocytes [36]. Accordingly, our *in silico* work introduces an interesting hypothesis into how VAR2CSA attaches more firmly *Pf* -infected erythrocytes to the proteoclycan matrix of the placenta when subjected to shear of the flowing blood. Overall, our computational results can be directly tested by experiments and propose, to our knowledge, a new mechanism of adherence of *Pf* -infected erythrocytes to the placenta. They may be relevant for the development of vaccines against placental malaria targeting the VAR2CSA protein.

## Author Contributions

CA-S designed the research. RR and NM carried out the simulations. All authors analyzed the data and wrote the paper.

## Acknowledgments

We thank Ulrich Schwarz, Motomu Tanaka, Michael Lanzer, and Manu Forero-Shelton for insightful discussions. We are grateful for financial support by the Klaus Tschira Foundation.

